# Influence of rifampicin-resistance mutations on the synthesis of antioxidant, DNA-protective, and SOS-inhibitory metabolites

**DOI:** 10.1101/2023.11.25.568633

**Authors:** V.N. Statsenko, E.V. Prazdnova

## Abstract

This work is aimed at studying the effects of rifampicin resistance mutations on the synthesis of secondary metabolites with antioxidant and DNA-protective properties.

We used probiotic strains of the genus *Bacillus*: *B. amyloliquefaciens* B-1895 and *B. subtilis* KATMIRA1933.

The antioxidant, DNA-protective activity, and the ability to suppress the SOS-response in *B. amyloliquefaciens* B-1895 and *B. subtilis* KATMIRA1933 rifampicin-resistant mutants have been studied for the first time. It has been found that antioxidant, DNA protective, and SOS-inhibiting activity is higher in rifampicin-resistant mutants than that of original strains.

According to the study results, it has been discovered that the antioxidant, DNA-protective and SOS-inhibitory activity in mutants *B. amyloliquefaciens* B-1895 and *B. subtilis* KATMIRA1933 resistant to rifampicin is higher than in control strains, which indirectly proves the pleiotropic effect of the *rpoB* gene on these activities.

## 1. Introduction

Many representatives of the genus *Bacillus* are known for their probiotic properties and are widely used in aquaculture (James et al, 2021), poultry farming (Popov et al, 2021), animal husbandry (Khalid et al, 2021) and medicine (WoldemariamYohannes et al, 2020).

*Bacilli* produce a wide range of secondary metabolites that provide antagonism to pathogens. Among the nonribosomal peptides (NRPs) produced by *Bacillus subtilis* and *B. amyloliquefaciens*, the most well-known are surfactin, which is an active biosurfactant that stimulates biofilm formation, and has antitumor, fungicidal, antibiotic activities and others (bacircin, lichenisin, pumilacidin) (Sharma, 2008; Farace, 2015). In addition, polymyxins, fusaricidins, iturins and fengycins with antibacterial and/or antifungal activity are of particular importance for agriculture (Ongena, 2008).

*B. subtilis* also produces lantibiotics, which are of a peptide nature and are formed through ribosomal synthesis. One of the most common lantibiotics is subtilin, which inhibits peptidoglycan synthesis and facilitates pore formation (Parisot, 2008). In addition to peptide antibiotics, *B. subtilis* and *B. amyloliquefaciens* can synthesize polyketides, which are synthesized by polyketide synthetases (PKSs) (Zhong, 2014). Thus, bacillin, dificidin, and macrolactin are synthesized with broad antibacterial activity against plant and human pathogens. Certain strains of *B. subtilis* synthesize polyene antibiotics with conjugated double bonds. Thus, hexaenes inhibit the growth of phytopathogenic fungi such as *Fusarium culmorum, F. sporotrichiella, F. oxysporum, Botrytis sorokiniana, Alternaria tenui*, and *Phytophthora infestans* (Sidorova, 2018).

*Bacillus* probiotics also generate agents that reduce the genotoxicity of substances such as 4-nitroquinoline-1-oxide, nitrosoguanidine, polyaromatic hydrocarbons, aflatoxins, and others (McBain, 2001, Lo, 2004). It has been found that metabolites of *B. amyloliquefaciens* B-1895 and *B. subtilis* KATMIRA1933 in cell-free preparations have DNA-protective activity and are able to suppress the SOS-response in *E. coli* (Mazanko, 2018). The mechanisms of such activity still require further study.

Suppression of the SOS response is of particular interest since it can also represent a mechanism of antagonism, but a more subtle one. It is known that SOS-induced mutagenesis can subsequently lead to the immunity of bacteria to antimicrobial drugs (Cirz, 2005). It has been shown that SOS-deficient bacterial strains adapt more slowly to antibiotics (Wigle, 2009). Moreover, activating the SOS-response by one therapeutic agent can lead to resistance to another agent (Chistyakov et al, 2018). A decrease in the intensity of the evolution of resistance may be one of the mechanisms of the fight against pathogens by probiotics.

At present, the search for antimutagens, as well as substances that can block the activity of genes and SOS-response factors, is an urgent task in pharmacology. Most of the synthetic inhibitors of SOS-repair are toxic (Alam et al, 2006), so it is necessary to look for natural bioavailable compounds.

Since rifampicin-resistant bacilli strains are known to produce more metabolites than wild-type strains (Cai, 2017), it was hypothesized that the production of DNA-protectors and SOS-inhibitors would be enhanced in rifampicin-resistant mutants of *B. amyloliquefaciens* B-1895 and *B. subtilis* KATMIRA1933.

## 2. Materials and methods

### 2.1. Cultivation of probiotic bacilli

We have used probiotic strains of the genus *Bacillus*: *B. amyloliquefaciens* B-1895 and *B. subtilis* KATMIRA1933 and Rif-mutants (Rif^R^), which were obtained from the collection of the Experimental Mutagenesis Laboratory in the Academy of Biology and Biotechnology, Southern Federal University. These strains are gram-positive soil bacteria and are used as probiotics in aquaculture and animal farming.

The cultivation of *Bacillus subtilis* KATMIRA1933 and *Bacillus amyloliquefaciens* B-1895 was carried out both in solid and liquid Luria-Bertani (LB) medium at 37 °C. The cell-free preparation used for testing was obtained by centrifuging the culture (Minispin-plus; Eppendorf) for 7 minutes at 6000 rps and filtration.

### 2.2. Obtaining Rif^R^ mutants

*B. amyloliquefaciens* B-1895 and *B. subtilis* KATMIRA1933 strains were inoculated on plates with LB medium with 0.04 mg/ml rifampicin. The cultivation of bacteria was carried out at 37 ° C for 3-4 days.

### 2.3. Biosensor strains

Recombinant strains of *E. coli* MG1655 (pRecAlux) and MG1655 (pKatG-lux) were used. The strains carry a recombinant plasmid with the LUX-operon of the luminescent bacterium *Photorhabdus luminescens*. This operon is transcriptionally fused with the promoters of the RecA and KatG protein genes, respectively. *E. coli* MG1655 biosensor (pRecA–Lux) emits light due to activation of the RecA promotor in response to DNA damage. This biosensor detects both nonspecific DNA damage (in the model with dioxidine) and the induction of the SOS-response by RecA activation (in the model with ciprofloxacin) (Chistyakov et al, 2018). The *E. coli* MG1655 (pKatG-lux) biosensor emits light due to activation of the pKatG promoter in response to oxidative stress (Zavilgelsky et al, 2007).

Recombinant strains of E. coli were grown in both liquid and solid LB medium with the addition of 100 μg/ml ampicillin at 37 ° C.

### 2.4. DNA damage and SOS-response inductors

Using the *Escherichia coli* MG 1655 (pRecA-lux) strain, the possibility of suppressing SOS-response and total DNA-protective activity was studied. 10-4 mg/ml ciprofloxacin and 2 · 10-5 mg/ml 1,4-dioxide 2,3-quinoxalinedimethanol (dioxidine) solutions in deionized water were used as inducers of SOS-response and DNA damage, respectively. *Escherichia coli* MG 1655 (pRecA-lux) strain was used to estimate antioxidant activity, with 10-3mg/ml hydrogen peroxide (Ferrain) solution in deionized water as an inducer.

Ciprofloxacin is a fluorinated quinolone with strong antibacterial activity. It is an antibacterial preparation with a wide range of action to which most gram-negative bacteria are very sensitive in vitro, and many gram-positive bacteria are susceptible or moderately susceptible. The primary mechanism of action of ciprofloxacin and other quinolones is to inhibit the bacterial DNA-gyrase, which plays a vital role in the process of matrix synthesis.

Dioxidine is an antibacterial preparation with a wide range of actions. By its chemical structure, dioxidine refers to the class of quinoxaline di-N-oxides. In vitro dioxidine activity varies depending on the type of bacteria, and this variation can be very large (Padeiskaya, 1977). It is assumed that the dioxidine action is based on suppressing the activity of bacterial deoxyribonuclease, which is the necessary enzyme of the reparation system of bacteria (Fadeeva, 1980).

### 2.5. Biosensor test

The biosensor test includes several stages:

1. Overnight cultivation of an *E. coli* biosensor strain.
2. Measurement of the density of night culture using a DEN-1B densitometer («Biosan») and standardization by adding LB liquid to 0.01 to McFarland.
3. Cell cultivation for 2 hours at 37°C to an early logarithmic phase.
4. 80 μl aliquots transfer to a 96-well plate. 10 μl of deionized water and 10 μl of the culture of the probiotic cell-free preparation are added to the control cells. In the experimental cells, 10 μl of the inductor (hydrogen peroxide, dioxidine, or ciprofloxacin) is added instead of water.
5. Bioluminescence measurement in a plate reader (FLUOstar Omega). Measurements are carried out every 10 minutes for 120 minutes.

Induction was evaluated by the formula:

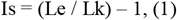

where (Is) - induction coefficient; Lk и Le - luminescence intensities of the control and experimental samples, respectively. A statistically significant excess of Le over Lk, assessed by the t-test, was considered as a sign of a significant influence on the induction effect. The protective effect (P, %) was calculated considering the induction in the presence of the corresponding tread concentrations according to the formula:

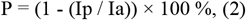

where Ip and Ia are the factors for the induction of the SOS response under the investigated influence in the presence of the protector and the control sample, respectively. All experiments were carried out in triplicate. Statistical analysis was performed using the Student’s t-test for P = 0.05.

## 3. Results

The antioxidant, DNA-protective and SOS-inhibiting activities of two strains have been studied in *B. subtilis* KATMIRA1933 and *B. amyloliquefaciens* B-1895, as well as Rif^R^ mutants of these strains. The antioxidant activity has been evaluated using a biosensor with the pKatG-lux construct. The DNA-protective and SOS-inhibiting activities have been evaluated using a biosensor with the pRecA–Lux construct.

Figure 1 presents the results of the study of the activity of the original and mutant strains of *B. subtilis* KATMIRA1933.

**Figure 1.**
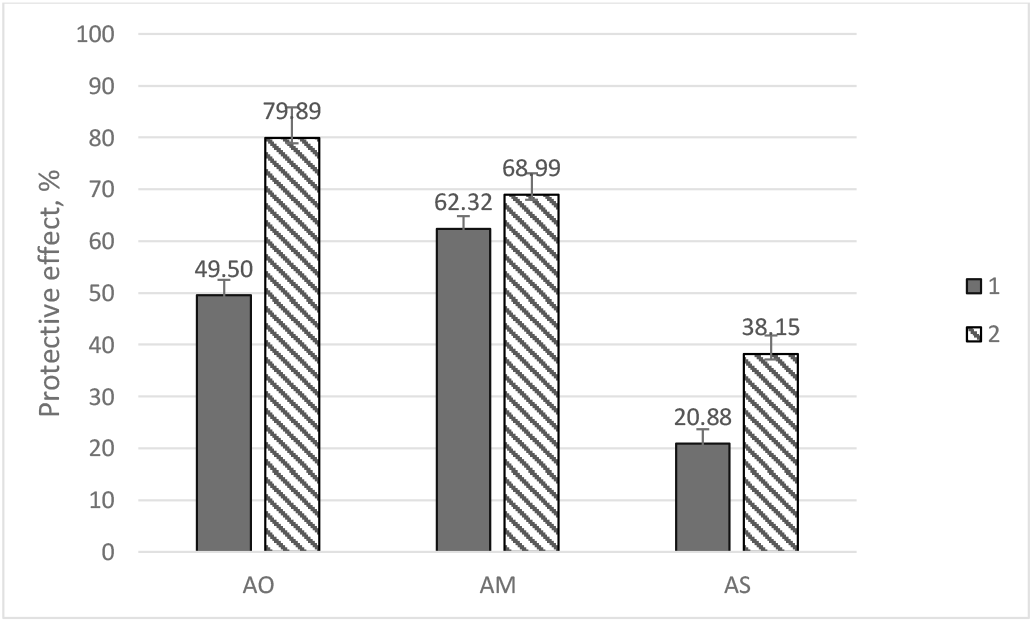
Protective activity of the original strain B. subtilis KATMIRA1933 and the Rifr mutant against hydrogen peroxide. AO – antioxidant, AM – antimutagenic (DNA-protective), AS – anti-SOS. 1-Original strain B. subtilis KATMIRA1933 in the presence of inducer; 2-Rifr mutant of B. subtilis KATMIRA1933 in the presence of inducer.

As can be seen from the presented data, the Rifr mutants have stronger antioxidant activity, which exceeds the original strain by 30,39 %. The DNA-protective activity of mutants also exceeds that of the original strain by 6,67%, and the ability to suppress the SOS response is higher by 17,27% in rifampicin resistant mutants. All differences are statistically significant, p<0.05.

For the second probiotic strain, quite similar data were obtained (Figure 2).

**Figure 2.**
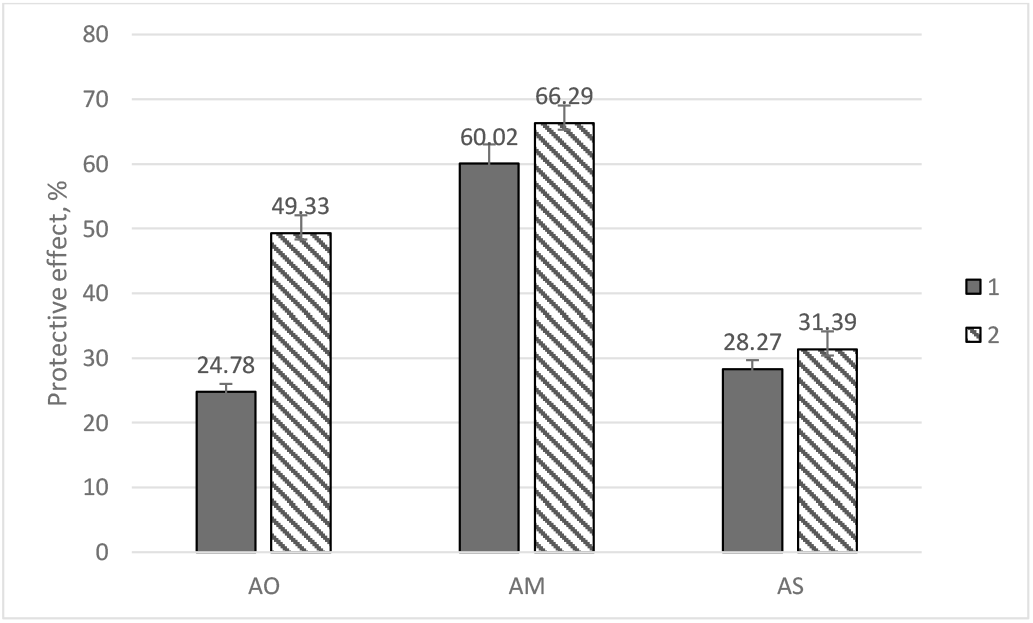
Protective activity of the original strain B. amyloliquefaciens B-1895 and the Rifr mutant against hydrogen peroxide. AO – antioxidant, AM – antimutagenic (DNA-protective), AS – anti-SOS. 1-Original strain B. amyloliquefaciens B-1895 in the presence of inducer; 2-Rifr mutant of B. amyloliquefaciens B-1895 in the presence of inducer.

As can be seen from the data above, all types of protective activity are stronger in the Rif^R^ mutants of *B. amyloliquefaciens* B-1895. The antioxidant activity of Rif^R^ strain metabolites is 24,55% higher than that of the original strain. The DNA-protective activity of mutant metabolites is 6,27% higher, and SOS-inhibitory activity is higher by 3,12% in rifampicin resistant mutants.

Comparing these two strains, one can say that both between the original strains and between Rif^R^ mutants, the metabolites of the *B. subtilis* KATMIRA1933 strain seem to be more active antioxidants than the metabolites of the *B. amyloliquefaciens* B-1895 strain. The antimutagenic and SOS-inhibitory activities of these two strains are close in range. It should be noted that the enhancement of SOS-inhibitory activity in mutants is much higher in the strain *B. subtilis* KATMIRA1933.

## 4. Discussion

Antioxidant and DNA -protector activities are shown for many types of probiotics, although bacilli are less commonly regarded as sources of antioxidants than, for example, lactic acid bacteria (Chooruk A., 2017). However, there are reports that representatives of the Bacillus genus can inactivate mutagens (McBain, 2001; Lo, 2004; Caldini et al, 2002; Genci et al, 2008; Jiang et al, 2016) and exhibit antioxidant properties (Wang et al, 2017; Rahman et al, 2018).

The SOS-inhibiting effect was previously demonstrated for metabolites of several probiotics (Chistyakov, 2018), however, the chemical nature of the substances that provide it is not yet established. The data we received indirectly indicates that this effect, like the DNA protector, can be provided by secondary metabolites.

It is known that rifampicin resistance arises due to mutations in the rpoB gene, which encodes the RNA polymerase (RNAP) β-subunit. Most Rifr mutations occur in a short (100 bp) area within rpoB. Rifampicin is associated with RNAP in a place adjacent to its active center, thereby physically blocking the formation of phosphodieter bonds in the main RNA circuit. Rifampicin inhibits any elongation of RNAs exceeding two or three nucleotides. These mutations reduce the affinity of RNAP for Rifampicin (Xu, 2005). Rifr mutations have pleiotropic phenotypes due to their transcriptional influence.

It has been found that rifampicin resistance mutations (in the *rpoB* gene) cause various physiological changes in bacteria, but such changes are not universal for all types of bacteria. A pleiotropic effect on rifampicin resistance is shown for other models - it is also known that B.velezensis Rifr mutants produce more secondary metabolites, in particular antifungal peptides of non-riboomal nature, such as iturin (Cai, 2017). In bacteria that are used to produce natural antibiotics, such as erythraezin (*Saccharopolyspora* e*rythraea*) and vancomycin (*Amycolatopsis orientalis*), specific *rpoB* mutations can increase the development of antibiotic data (*Koch*, 2014). Rifampicin-resistant *Pseudomonas protegens* mutants, which carry H531n in the -subunit RNA polymerase, show improved antifungal activity because of the synthesis of two antibiotics: 2,4-diacetyloflusion (Phl) and pivotine (Plt) (X*ie*, 2016). In *Escherichia coli* carrying a mutation with a frameshift mutation in the *rpoB* gene, the level of production of protein metabolites in mutants is higher than that of wild type, which is due to the feedback mechanism (Huseby, 2020). It has been found that the biosynthesis of actinochordin (ACT), undecilprodigiosine (Red), and calcium-dependent antibiotic (CDA) is activated by introducing mutations to the *rpoB* gene in *Streptomyces lividans*, which emits less or does not produce antibiotics at normal conditions. It has been shown that the missense mutation H437Y in the *rpoB* gene activates anthrahamicin biosynthesis in *Streptomyces chattanogensis* compared to the wild type (L*i*, 2019). It has also been found that Rif^r^ mutants of *Streptomyces diastatochromogenes* show increased production of the nucleoside antibiotic tooamicine (TM). A mutant carrying an amino acid substitution of HIS437Arg was the best producer of TM, with an increase of 4.5 times compared to a wild-type strain (Ma, 2016).

Thus, the obtained data indirectly confirm the hypothesis that the SOS-inhibiting and antimutagenic activities, including the antioxidant activity of probiotics, can be provided by secondary metabolites, non-ribosomal peptides, since their products increase in Rif^R^ *Bacillus* mutants.

The strains used are promising probiotics, which are used both in aquaculture and agriculture. *B. amyloliquefaciens* B-1895 is used as a probiotic and a replacement for antibiotics in aquaculture for growing *Alburnus leobergi* (Karlyshev, 2014). *B. subtilis* KATMIRA1933, when used as a feed additive, has shown multiple beneficial effects on chickens (Chistyakov, 2018).

Therefore, it seems reasonable to look for ways to increase the possibility of the excretion of secondary metabolites by these strains. Strategies for producing more effective probiotic strains may include a selection of spontaneous mutants or induced mutagenesis targeting genes with pleiotropic action.

It should be borne in mind, though, that rifampicin is widely used to treat tuberculosis and other socially significant diseases (Grobbelaar M. et al., 2019), so the spread of resistance to it should be limited. Therefore, it would be unwise to use Rif^R^ strains as probiotics, as they may serve as a source of genes for antibiotic resistance in the host microbiome due to horizontal gene transfer. However, they can be used as producers of cell-free preparations of antioxidants, DNA protectors, and SOS-inhibitors.

## 5. Conclusion

Thus, it has been found that antioxidant, DNA protective, and SOS-inhibiting activity is higher in rifampicin-resistant mutants of *B. subtilis* KATMIRA1933 and *B. amyloliquefaciens* B-1895 than that of original strains. The data obtained indirectly confirms the possibility of non-ribosomal metabolites participating in these activities. The results of the work can be applied as the basis for creating strains that are producers of secondary metabolites with the above-described biological activities.

## Data availability statement

Data generated or analyzed during this study are provided in full within the published article.

